# FAKHRAVAC and BBIBP-CorV vaccine seeds’ binding to angiotensin-converting enzyme 2: A comparative molecular dynamics study

**DOI:** 10.1101/2023.10.19.563051

**Authors:** Soroush Setareh, Iman Rad, Jafar Meghdadi, Kaveh Khodayari, Ahmad Karimi Rahjerdi

## Abstract

**Background:** Safety and efficacy of the SARS-CoV-2 inactivated vaccines have been question since the emergence of SARS-CoV-2 variants of concern (VOCs). Using residue fluctuations and statistically comparing RMSF values, have escalated the understanding of the binding dynamics of the viral proteins to their receptors and here in this study, we compared the interaction between inactivated spike proteins (representing FAKHRAVAC and BBIBP-CorV vaccines seed) and the human Angiotensin-Converting Enzyme 2 (hACE2) receptor.

**Methodology:** Through 100 set of accelerated 1 ns comparative molecular dynamics simulations, we analyze the binding dynamics and energy components of these interactions and compared residue backbone fluctuations using entropy and statistics including KL-Divergence and KS-test.

**Principal Findings:** Our results reveal that FAKHRAVAC and Sinopharm exhibit similar binding dynamics and affinity to hACE2. Further examination of residue-wise fluctuations highlights the common behavior of binding key residues and mutation sites between the two vaccines. However, subtle differences in residue fluctuations, especially at critical sites like Q24, Y435, L455, S477, Y505, and F486, raise the possibility of distinct efficacy profiles.

**Conclusion:** These variations may influence vaccine immunogenicity and safety in response to evolving SARS-CoV-2 variants. The study underscores the importance of considering residue-wise fluctuations for understanding vaccine-pathogen interactions and their implications for vaccine design.

**Author summary:** It is fundamentally important to ensure the safety and efficacy of the FAKHRAVAC, as an inactivated vaccine candidate for SARS-CoV-2. Considering the previously published pre-clinical and clinical findings about the similarity of the FAKHRAVAC’s safety and efficacy in comparison to the BBIBP-CorV vaccine seed (which is recalled as Sinopharm), it is necessary to gain more insights into structure and function of this vaccine at the molecular level, as well. Since the binding dynamics of the viral proteins to their receptor can imply the vaccine’s immunogenicity and mechanism-of-action, binding dynamics of a vaccine candidate must be studied comprehensively. Hereby, we have compared binding dynamics of the FAKHRAVAC and Sinopharm vaccine seeds to the SARS-CoV-2 spike protein’s receptor, the ACE2. We took advantage of a comparative molecular dynamics simulation approach to effectively compare binding dynamics using atom fluctuations and at the residue level to ensure the resolution of this study. We have found similar binding dynamics and binding mechanics between these two vaccines, validating the pre-clinical and clinical findings computationally, as well as highlighting residues with different fluctuations and discussed their potential roles.

## 1- Introduction

The rapid emergence of SARS-CoV-2 (Severe Acute Respiratory Syndrome Corona Virus 2) variants of concern (VOCs) has raised concerns regarding fast and intelligent vaccine development. In the case of the inactivated vaccines, which were based on the wild-type spike (S) protein [1, 2], it was doubtful whether they could provide sufficient protection against the emerging antigenically distinct VOCs [3]. Since VOCs refer to variants, which have been demonstrated clinically, such as high disease transmissibility and severity, and decreased neutralization antibody developed against previous infection or vaccination [4], a serious concern has emerged regarding how amino acid substitutions in the receptor-binding domain (RBD) of the virus’s spike glycoprotein have something to do with the main region involved in ACE2 binding and viral pathogenicity [5].

Several VOCs have been introduced since December 2020, such as B.1.1.7 (Alpha), B.1.351 (Beta), etc. [4, 6], which all obtained mutations regarding the spike protein strain that was originally found in Wuhan [4]. In this study, the interaction of inactivated Spike proteins (*i.e.* the antigenic component of FAKHRAVAC [7] and Sinopharm [8] vaccines’ viral seed) with a target protein (*i.e.* human Angiotensin-Converting Enzyme 2: hACE2) [3] was simulated. Considering that both inactivated vaccines have confirmed published clinical trial results [9, 10] with rather similar evaluation indices, determining the correlation between the mechanism of protein– protein interaction and either clinical or para-clinical manifestation was the aim of the current study.

## 2- Methods and materials

### 2-1- Protein Structures and Preprocessing

The three-dimensional (3D) structure models for Spike proteins corresponding to Sinopharm were obtained from the CryoEM structure of the SARS-CoV-2 Spike protein (PDB ID: 7N9T). The 3D structures of the human angiotensin-converting enzyme 2 (hACE2) receptor were obtained from the Protein Data Bank (PDB ID: 7L7F).

The SARS-Cov2 nucleotide sequence that was acquired from the patient’s sample as the seed of the FAKHRAVAC vaccine was sequenced using Illumina Mi-Seq. The obtained viral genome sequence was then transcribed into its corresponding amino acid residues. Then, the 3D structure was determined on the basis of the 7N9T template using Molegro virtual docker, as explained in previous studies [11, 12]. The FAKHRAVAC structure contains two major amino acid substitutions compared to its homolog template (*i.e.* 7N9T) used in the model, which are D138Y and S477N residues.

All protein structures were preprocessed before further analysis. Bonding hydrogens were added, and crystallographic water molecules were removed.

### 2-2- Protein docking and molecular dynamics simulation

Binding sites are the interface between Spike proteins and their receptor (hACE2), which introduce active and passive residues. Regarding active residues located in the binding interface, protein docking was performed using the High Ambiguity Driven protein– protein DOCKing (HADDOCK 2.4) webserver [13]. Spike proteins corresponding to FAKHRAVAC and Sinopharm were docked with the target protein hACE2.

Docked structures then undergo cleaning to prepare the simulation system. A Python script is used to clean the complex. The GROMACS topology for all complex structures was written using the GROMACS 2022.2 package [14]. CHARMM-36 (March 2019 version) is used throughout all simulations as the forcefield. For each system, complexes were placed in 125 x 125 x 125 Å dodecahedron boxes (volume of 1,953 nm^3^) solvated in SPC/E 216 water molecules, as explained previously [15]. The system charge was then neutralized by substituting solvent molecules into positive (Na^+^) and negative (Cl^-^) counter-ions.

### 2-3- Comparative molecular dynamics (MD) analysis

The comparative MD simulations were performed through 100 sets of 1-ns short MD simulations, as described by *Rajendran et al.* [16]. In brief, simulation systems including complexes and solvent atoms are subjected to energy minimization, where the system potential energy is minimized using the steepest decent minimization algorithm with the selection of the particle mesh Ewald method as the treatment of long-range electrostatic interactions. Followed by equilibration for a 300 ps process at 300 K and ambient pressure. Then, a 10-ns equilibration proceeded. The resulting simulation is then extended for a random period of 0–0.5 ns for each replication set to ensure the natural randomness of motions and randomization of the initial state. Each replication set is then simulated for 1 ns.

### 2-4- Binding free energy analysis

Binding free energy (BFE) for the process of binding to hACE2 was analyzed for both vaccine spike proteins. Calculation of BFE and its decomposition into energy terms is performed using the GMXMMPBSA library as implemented in the bash environment. GMXMMPBSA uses the Generalized Born equation for the calculation of energy terms as denoted in equation (1):

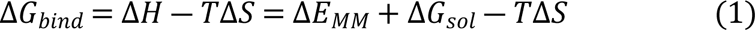

Where the Δ*E*_*MM*_ and Δ*G*_*sol*_ denote molecular mechanics and solvation free energy, respectively, and can be expanded further into energy terms as described in Equations (2) and (3):

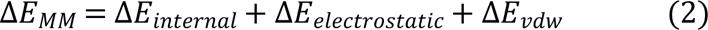

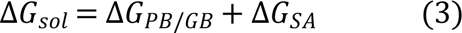

Where Δ*G*_*PB*/*GB*_ and Δ*G*_*SA*_ are showing Poisson-Boltzmann (Generalized Born) and solvent-accessible surface area free energies, respectively. The energy terms are then compared to each other using a two-sampled t-test.

### 2-5- Analysis of MD Results

To ensure that the simulation systems are equilibrated, backbone root mean square deviation (RMSD) is calculated for each protein in both complexes.

To effectively compare the process of binding of the spike protein to ACE2, root mean square fluctuation (RMSF) is calculated for each residue of both spike protein structures. RMSF is calculated as equation (4) denotes:

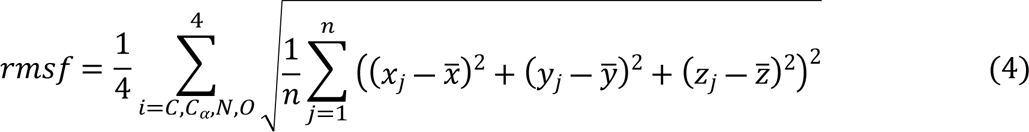

where *x,y,* and *z* are the coordinates of a given atom through j timepoints and the 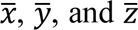 are the average coordinates of that atom through all timepoints. The RMSF is calculated for the backbone of each residue, as is calculated for each of C, C_α_, N, and O atoms, and is the averaged over the backbone atoms of each residue. To compare the RMSF values between the residues of each complex, entropy (which should not get confused with the Entropy as is in the physical chemistry) is used, as defined in the KL divergence formula (5):

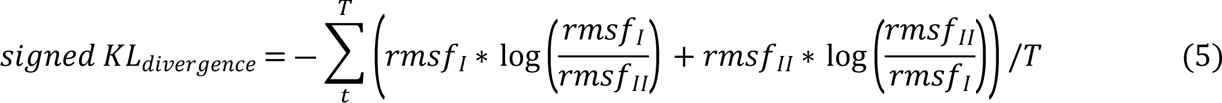

where rmsf_I_ and rmsf_II_ are the rmsf of FAKHRAVAC and Sinopharm, respectively, both in the bound state to hACE2. The rmsf values are treated as probability vectors and normalized using min– max normalization, as denoted in equation (6). The KL_divergence_ is finally signed to maintain the definition of relative entropy.

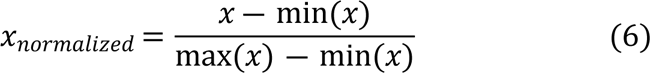

### 2-6- Identification of key residues

The two-sample KS (Kolmogorov-Smirnov) test is then used to statistically compare RMSF values over all replicates. The *p-*values correspond to the KS test, resulting in *D*-values, which are then corrected considering the false discovery rate by the Benjamini– Hochberg method [17]. Residues with significant *p-*values were then identified.

## 3- Results

Site-wise comparison of atom motions between two dynamical states results in the identification of residues with dampened motion, both in the Spike protein and ACE2 receptor structures. We have investigated both residues with and without change in the fluctuations between two complexes and have also tested the significance of the difference, since they might impose an important role, either structural or functional. The first set of results represents four complexes of interest that were equilibrated prior to the comparative accelerated MD simulations (Fig 1).

**Figure 1.**
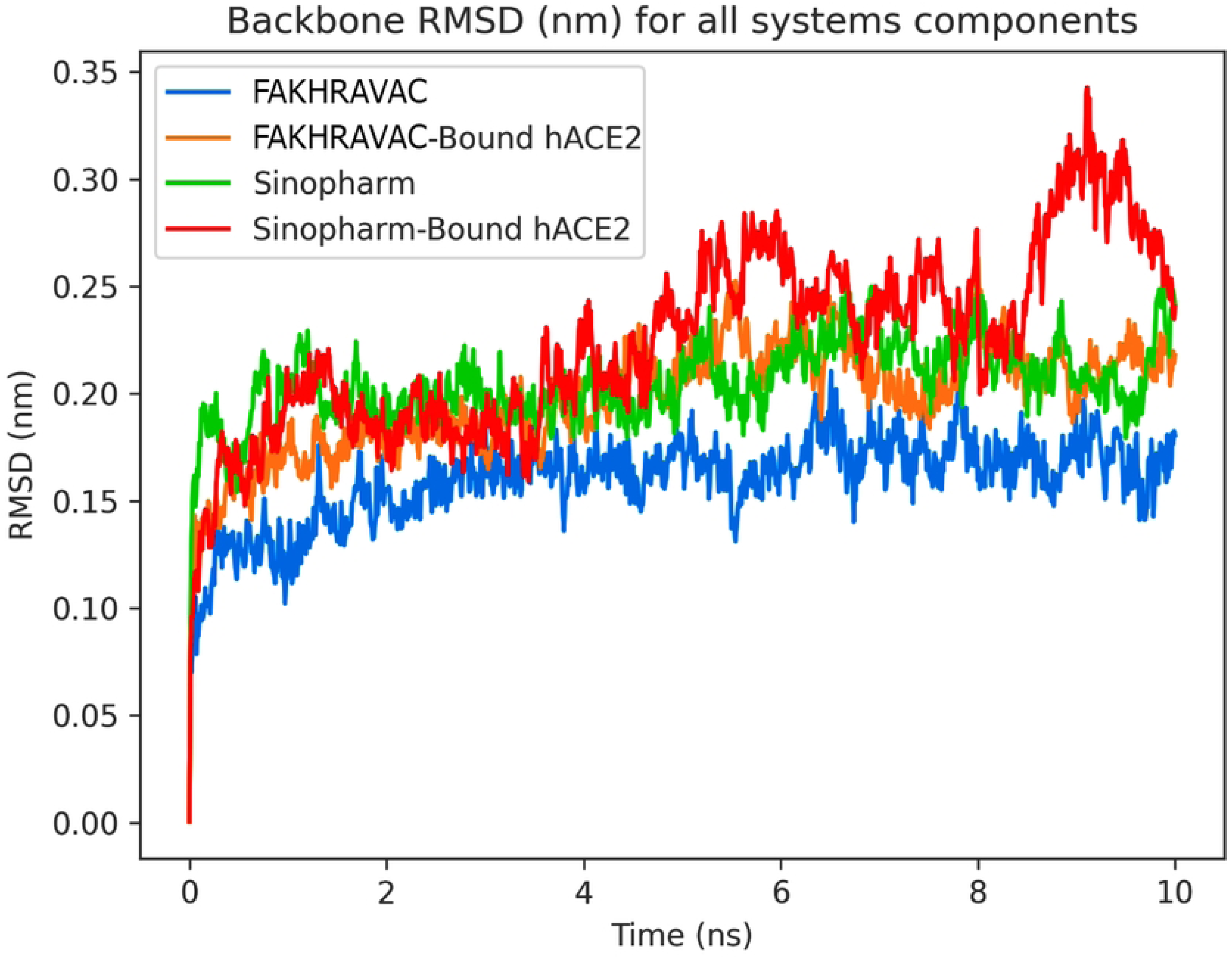
Backbone RMSD plots of the equilibration simulation. Backbone RMSD plots for FAKHRAVAC (Blue), Sinopharm (Green), and hACE2 in FAKHRAVAC-bound (Orange) and Sinopharm-bound (Red) states.

### 3-1- FAKHRAVAC and Sinopharm bind to hACE2 at the same energies

Total binding energy determines the affinity of the spike protein toward its receptor, the, and the kinetics and thermodynamics of subsequent molecular and cellular events, including viral entry into cells, which directly reflects the transmissibility and immunogenicity of the viral particles. Because different residue interactions can impose different binding energies, both in terms of binding energy compartment and binding energy value, it is important to analyze energy compartments solely, in addition to considering total binding energy.

Binding free energy (BFE) decomposition to BFE energetic components facilitates comparison of binding thermodynamics between the two vaccines, which are found to be similar. Despite a similar BFE contribution, FAKHRAVAC shows a slightly stronger electrostatic binding energy than Sinopharm. However, despite the more unfavorable solvation of the FAKHRAVAC structure, the binding energy of both vaccines remains the same (Fig 2).

**Figure 2.**
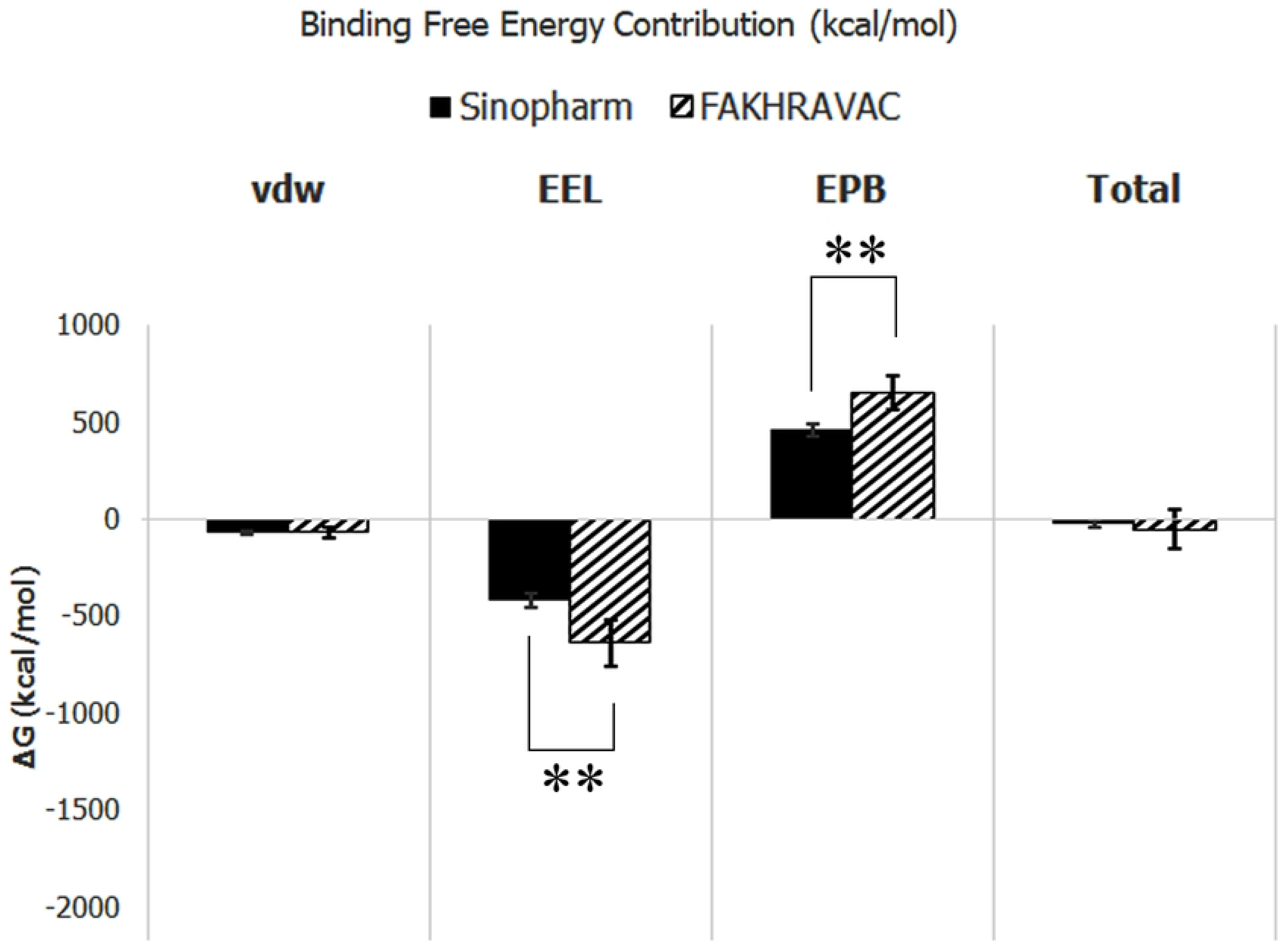
Spike protein binding to hACE2and contribution to binding free energy (BFE). The vdw (van-der-Waals energy), EEL (Electrostatic energy), EPB (Poisson-Boltzmann energy), and total indices of BFE in FAKHRAVAC and Sinopharm are shown in black and dashed series, respectively.

Electrostatic energy and Poisson– Boltzmann energy were significantly different between two vaccines with *p*-values < 0.05, but van-der-Waals energy was similar. In total, BFE of the two vaccines binding to hACE2 remains the same with no significant change.

### 3-2- Altered atom motions between FAKHRAVAC and Sinopharm

The site-wise comparison of the RMSF values during all 100 sets of short MD simulations shows which residues fluctuate significantly different from the corresponding residue in the other complex structure. The Kolmogorov– Smirnov statistical test (KS-test) is applied to test the null hypothesis of equal RMSF values. Furthermore, Fig 3-a represents which residues have significantly different sets of fluctuations.

**Figure 3.**
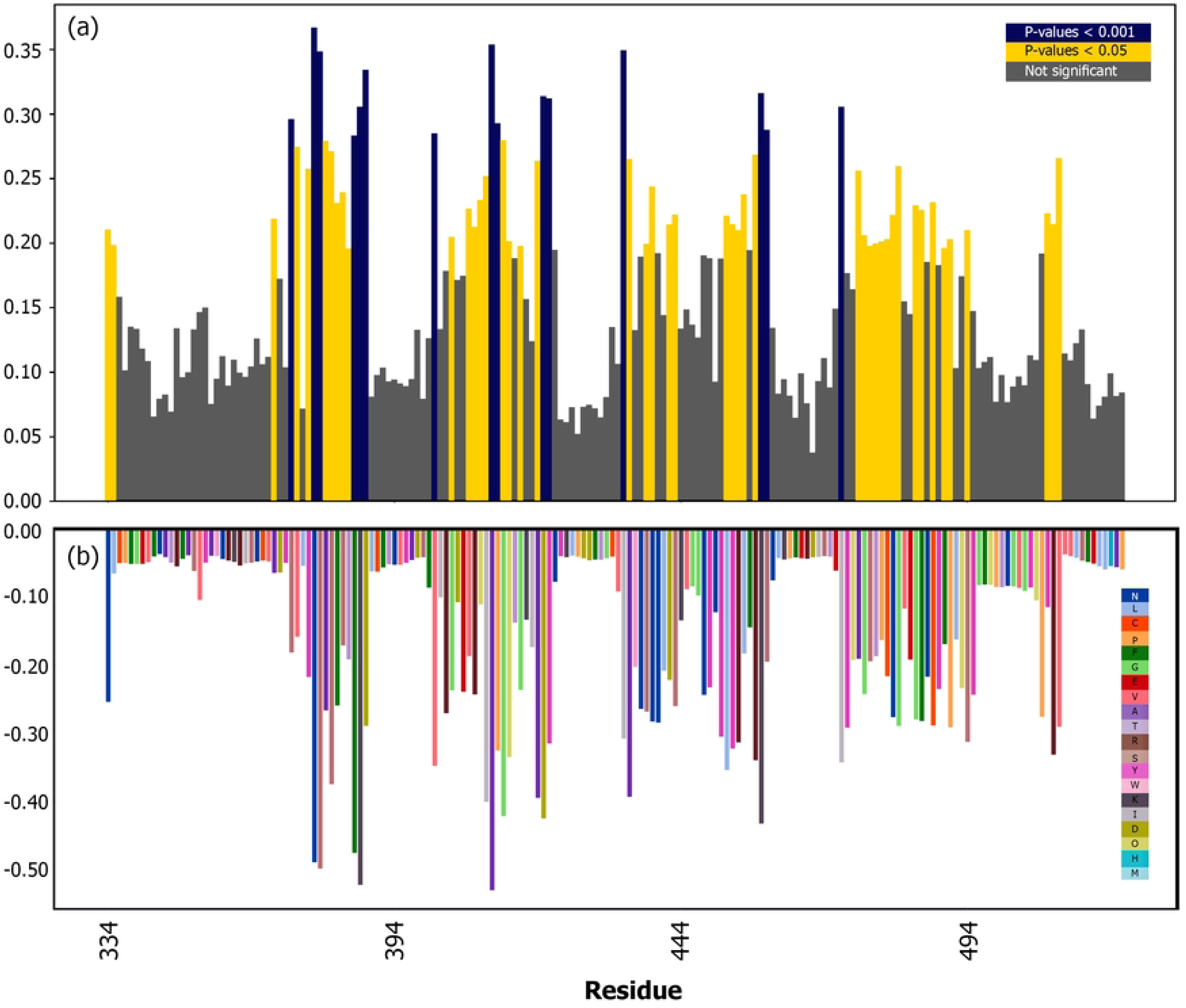
Comparative backbone RMSF of FAKHRAVAC and Sinopharm residues. (a) The strongly significant (p-value < 0.001) and significant (p-value < 0.05) residues are colored blue and yellow, respectively. (b) Signed KL divergence (dFLUX) between FAKHRAVAC and Sinopharm states bound to human ACE2, with color-coded residues.

Here, we also compared two distributions of RMSF values, one corresponding to the FAKHRAVAC motions and the other corresponding to the Sinopharm motions. The RMSF distributions are assumed to be probability distributions and then compared to each other using the Kullback– Leibler divergence (KL-Divergence) metric, which was originally used to compare probability distributions. In this regard, the more negative the signed KL-Divergence the more distinct the two probability distributions. In other words, KL divergence here indicates how surprising the Sinopharm RMSF values would be, given the probability distribution of FAKHRAVAC RMSF.

Negative dFLUX (signed KL-Divergence) values indicate a shift in the RMSF distributions between two defined dynamic states, FAKHRAVAC and Sinopharm. Residues with negative dFLUX values, as shown in Fig 3-b, show different fluctuations between FAKHRAVAC and Sinopharm, suggesting different behaviors and binding mechanisms in those residues.

KL divergence is a directionless metric not sensitive to using the states interchangeably, where negative values of dFLUX indicate dampened atom motions, which can be simplified as only atom motion alteration.

### 3-3- Binding mechanics of inactivated vaccines with human ACE2

Atom motions between the two dynamic states of ACE2 binding, Bound to FakhraVac and Bound to Sinopharm, are also compared, both as probability distributions (Fig 4-a) and by comparing the RMSF values themselves (Fig 4-b).

**Figure 4.**
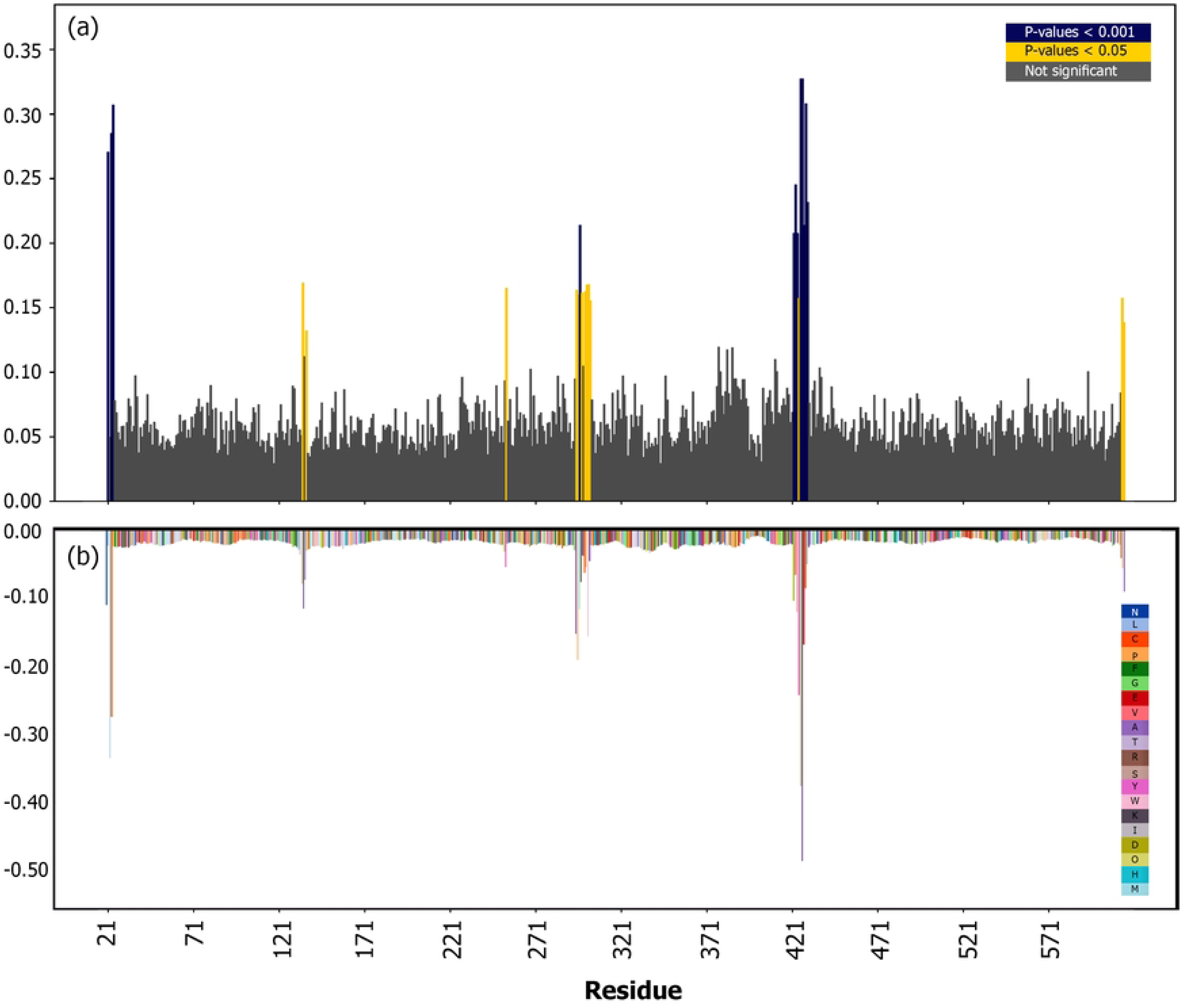
Comparative RMSF of the ACE2 backbone residues when comparing FAKHRAVAC-bound and Sinopharm-bound dynamic states. Strongly significant (p-value < 0.001) and significant (p-value < 0.05) residues are colored as (a) blue and yellow, respectively. (b) Signed KL divergence (dFLUX) of ACE2 between FAKHRAVAC-bound and Sinopharm-bound dynamic states, with color-coded residues.

The dFLUX values are illustrated as a heatmap over hACE2 and Spike protein residues, showing altered atom motions (Figure 5). As can be seen from the dFLUX heatmap, most of the hACE2 residues show no change in the fluctuations between FAKHRAVAC and Sinopharm, suggesting similar binding mechanics between the two vaccines and their receptors.

**Figure 5.**
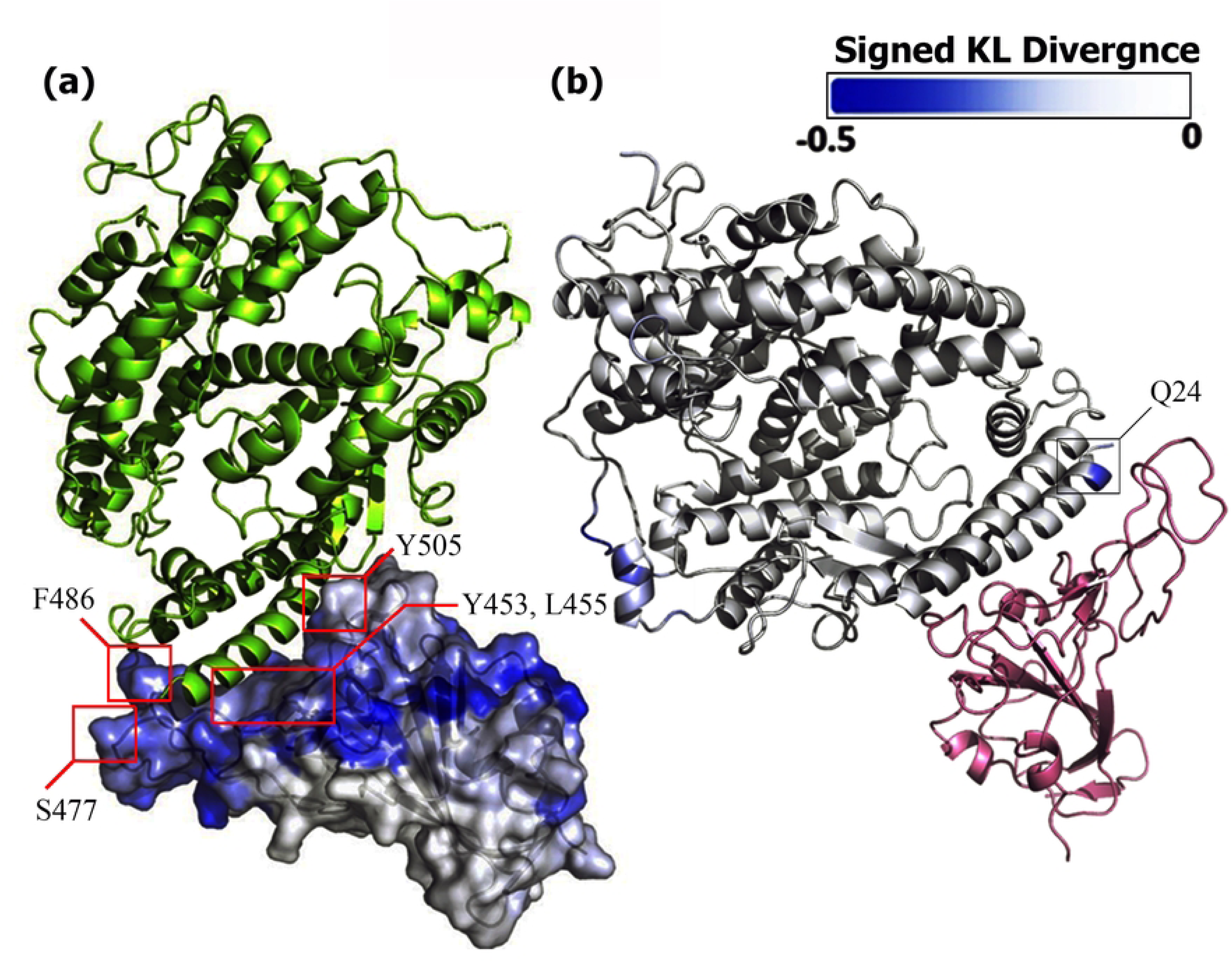
hACE2 and spike protein complex with a signed KL-divergence gradient over each protein’s residues. (a) Signed KL-divergence values are demonstrated over the spike protein’s surface, and residues with significant KS-test *p*-values and previously indicated in the literature are outlined in red. (b) Signed KL-divergence values are shown for the hACE2 residue, where the Q24 residue is outlined.

Despite the KL divergence being the same in nearly all residues of hACE2, some residues also show different fluctuations when facing other vaccines. The I23, E23, Q24, M297, G422, L423, L424, P425, D427, F428, Q429, and E430 residues show strongly significant (*p*-value < 0.001) and residues P135, N137, S254, D295, A296, V298, Q300, A301, W302, D303, and S425 show significant (*p*-value < 0.05) differences in fluctuations when bound to FAKHRAVAC compared to the state bound to Sinopharm.

Even spike proteins that are very similar in structure still show different KL divergence values, which indicates different binding mechanics.

## 4- Discussion and conclusion

The results of the current study indicate that despite the mostly significant fluctuations in the spike proteins, the motion of the residues located at the binding interface and binding key residues have either remained the same or have changed slightly, which are not significant to describe an alternated binding dynamic between FAKHRAVAC and Sinopharm. To effectively compare residue fluctuations, we observed the behavior of the FAKHRAVAC and Sinopharm spike proteins and the ACE2 receptor through a set of short MD simulations, in addition to considering the randomness and chaotic nature of the initial protein configuration. Then, the observations of protein fluctuations are compared statistically.

### 4-1- Binding Dynamics and Residue Fluctuations

The pattern of how fluctuations change through binding processes provides valuable insights into the dynamics of binding. The process of spike protein binding to the ACE2 receptor itself generally reduces RMSF [16]; however, the dampened motion profile differs from one virus seed to another. Although similar fluctuations suggest similar binding dynamics in general, it should be noted that even minor differences might play an important role in amino acid interactions when facing different binding processes. Both in the spike protein and ACE2 structures, a number of residues are found with significantly different fluctuations, which are neither the residues involved in the binding process nor the binding key residues. This finding suggests that the motion of residues involved in the binding process is the same between the two vaccines. On the other hand, other binding key residues and mutation hotspots, specifically mutation sites between major SARS-CoV-2 strains or residues predicted to be pathogenic, fluctuate the same either in FAKHRAVAC or Sinopharm. These findings indicate similar binding mechanics and hence, probably similar immunogenicity and safety profiles, as also shown in FAKHRAVAC pre-clinical [7] and clinical studies [2], confirming and computationally validating previous clinical findings.

### 4-2- Distinct Residue Fluctuations

Despite mostly similar binding dynamics of FAKHRAVAC and Sinopharm, some residues do fluctuate significantly both on the spike protein’s surface and on the receptor’s surface. Some of these residues are either harboring mutations between different SARS-CoV-2 strains or are classified as key binding residues. Binding key residues identified in previous studies and residues with significantly different fluctuations found in our study are listed in Table 1. As shown in Table 1, where the interacting residues are identified mutually, they are highlighted in bold.

**Table 1.**
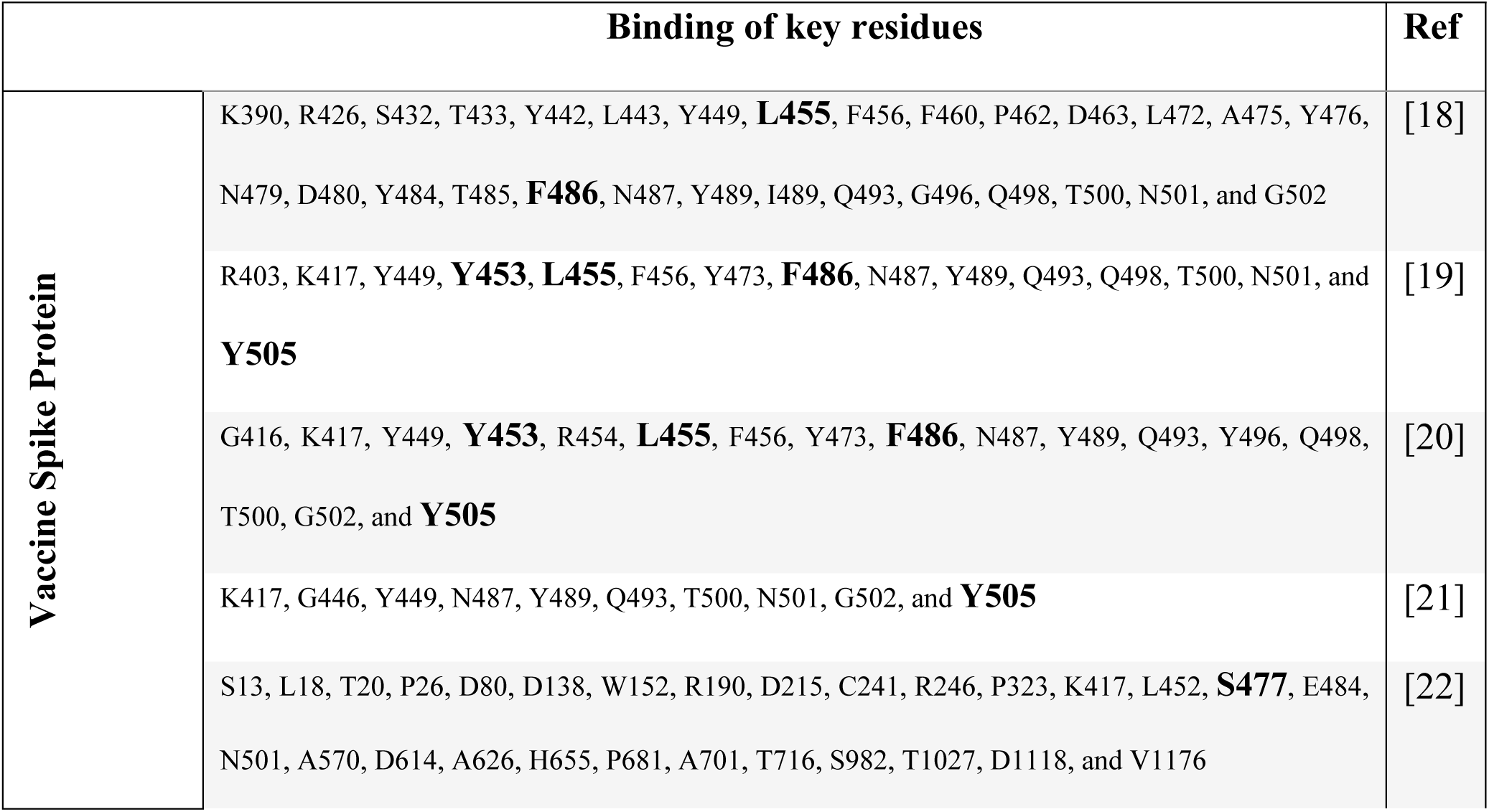
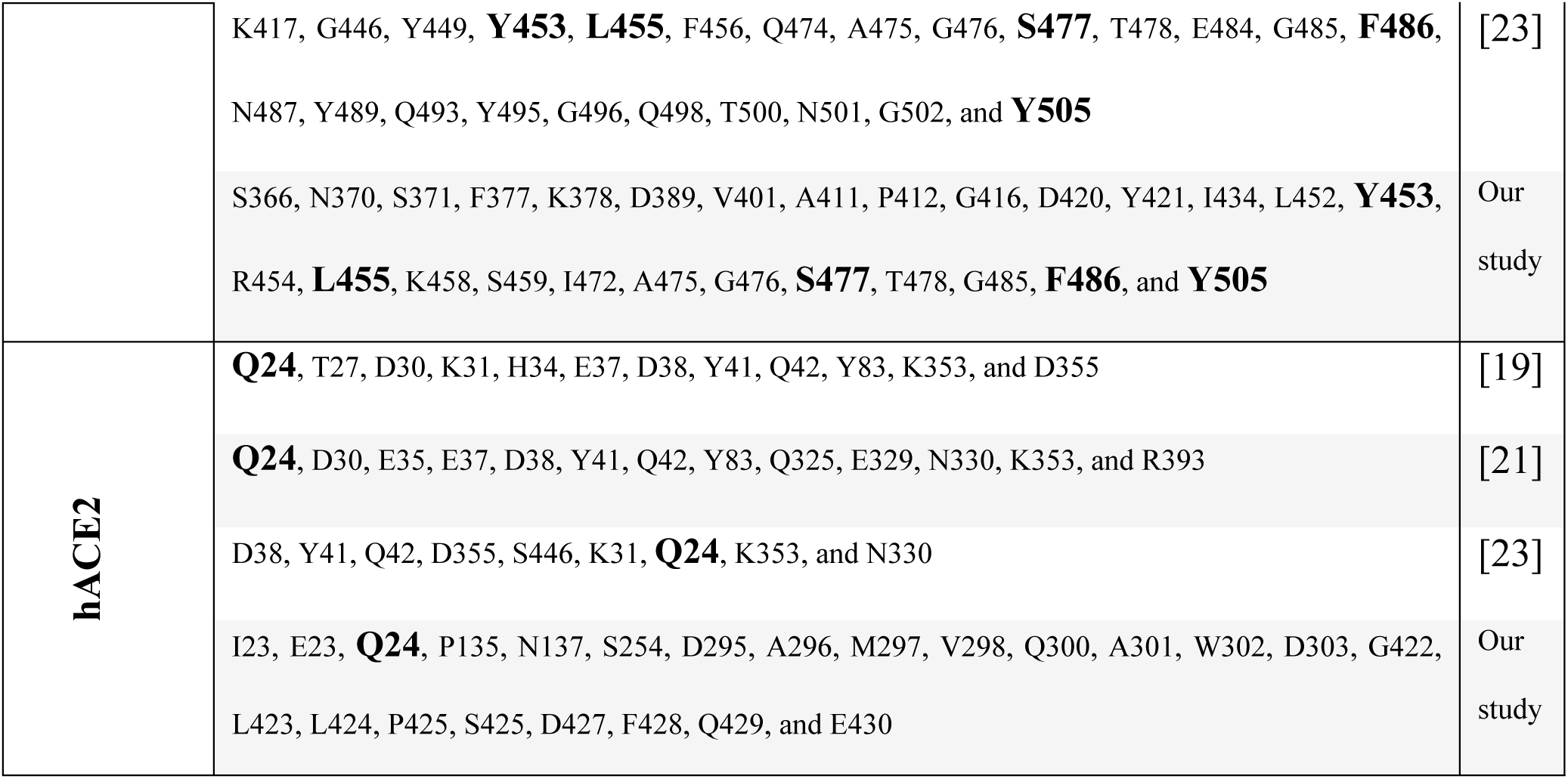
Binding key residues identified through either previous studies or residues with significantly different fluctuations between FAKHRAVAC-hACE2 and Sinopharm-hACE2 complexes.

| Binding of key residues |  | Ref |
| --- | --- | --- |
| Vaccine Spike Protein | K390, R426, S432, T433, Y442, L443, Y449, **L455**, F456, F460, P462, D463, L472, A475, Y476, N479, D480, Y484, T485, **F486**, N487, Y489, I489, Q493, G496, Q498, T500, N501, and G502 | [18] |
|  | R403, K417, Y449, **Y453**, **L455**, F456, Y473, **F486**, N487, Y489, Q493, Q498, T500, N501, and **Y505** | [19] |
|  | G416, K417, Y449, **Y453**, R454, **L455**, F456, Y473, **F486**, N487, Y489, Q493, Y496, Q498, T500, G502, and **Y505** | [20] |
|  | K417, G446, Y449, N487, Y489, Q493, T500, N501, G502, and **Y505** | [21] |
|  | S13, L18, T20, P26, D80, D138, W152, R190, D215, C241, R246, P323, K417, L452, **S477**, E484, N501, A570, D614, A626, H655, P681, A701, T716, S982, T1027, D1118, and V1176 | [22] |
|  | K417, G446, Y449, **Y453**, **L455**, F456, Q474, A475, G476, **S477**, T478, E484, G485, **F486**,<br>N487, Y489, Q493, Y495, G496, Q498, T500, N501, G502, and **Y505** | [23] |
|  | S366, N370, S371, F377, K378, D389, V401, A411, P412, G416, D420, Y421, I434, L452, **Y453**,<br>R454, **L455**, K458, S459, I472, A475, G476, **S477**, T478, G485, **F486**, and **Y505** | Our<br>study |
| hACE2 | **Q24**, T27, D30, K31, H34, E37, D38, Y41, Q42, Y83, K353, and D355 | [19] |
|  | **Q24**, D30, E35, E37, D38, Y41, Q42, Y83, Q325, E329, N330, K353, and R393 | [21] |
|  | D38, Y41, Q42, D355, S446, K31, **Q24**, K353, and N330 | [23] |
|  | I23, E23, **Q24**, P135, N137, S254, D295, A296, M297, V298, Q300, A301, W302, D303, G422,<br>L423, L424, P425, S425, D427, F428, Q429, and E430 | Our<br>study |

Because different fluctuations indicate different roles in the binding process, we can anticipate that the differences in the binding dynamics are rooted in the fluctuations of these residues. Even though they tend not to show a significant binding alteration between the two vaccines in this study, it might highlight some clues about possible separations between FAKHRAVAC and Sinopharm efficacy in the future, regarding the occurrence of variations in these specific residues.

Theoretically, any direct (at the same residue level) or indirect (at the spatially adjacent residues) structural variation might change the interactions and atom fluctuations. Therefore, altered atom fluctuations at the site of residues presented in this paper can become potentially intolerable through the interaction of ACE2 and spike proteins creating different binding dynamics and eventually different immunogenicity and safety profiles of the vaccine.

Different residue fluctuations, dampened atom motions, and rigidity of the RBD– ACE2 complex. Furthermore, it determined how strongly the spike protein and its receptor were bound together, mirroring the affinity to the receptor. Reductions in the RMSF values and dampened atom motions are shown through the simulation of different SARS-CoV-2 VOCs, including the Omicron.

#### 4-2-1- hACE2 Residue Fluctuations

The Q24 residue on the surface of ACE2, for instance, shows significantly different fluctuations when bound to FAKHRAVAC compared to the state bound to Sinopharm. Q24 has been continuously reported in the literature as one of the most critical residues of hACE2 in antibody interactions, formation of hydrogen bonds, and other interactions [24-26].

#### 4-2-2- Spike Protein Residue Fluctuations

The Y435 residue is one of the spike protein’s mutation sites [27], capable of forming hydrogen bonds and priming molecular interactions when the D435Y mutation occurs [28, 29], which is also a focus of vaccine design studies [30]. The L455 residue is a key residue in binding to hACE2, neutralizing antibody escape, and forming molecular interactions. [31-35]. The S477 residue is one of the other dangerous residues in the spike protein structure, which corresponds to receptor binding and neutralizing antibody escape [32] and is a mutation site formed under selective pressure and contributing to higher infectivity and immune escape of the Omicron variant [36-38]. The role of S477N substitution and its impact on immune escape and its molecular and clinical consequences has been well discussed previously [39]. The Y505 and F486 residues are also key mutation sites, corresponding to stronger interactions with hACE2 [40-42], and are also key binding residues [43, 44]. The F486L mutation is highly infectious and resistant to vaccine sera, highlighting the role of this residue in immune escape and viral internalization [45].

The flexibility of the protein residues can imply protein function and mobility [46]. In the case of viral proteins, RMSF can describe virus pathogenicity. Higher fluctuations could be interpreted either as looser binding and lower affinity or the viral protein’s ability to adapt itself to different receptor structures [47]. The impact of changing residues on the flexibility of SARS Coronavirus proteins through mutations and the resulting different binding affinities and mechanics has been shown by a variety of prior studies. Considering the vaccine mechanism of action and spike protein-facilitated entry of viral particles into target cells, comprehensive study of the protein regarding its residue-wise fluctuations, both at the ligand and receptor scales, is required.

### 4-3- Future Research Directions

This study aimed to evaluate the spike protein’s affinity for its receptor, hACE2, in the context of FAKHRAVAC, using computational biology and molecular dynamics. Future research should address the interaction between the vaccines and other immune components, particularly spike neutralizing antibodies, to fully understand their immunogenicity.

